# Mice lacking *Ptprd* exhibit deficits in goal-directed behavior and female-specific impairments in sensorimotor gating

**DOI:** 10.1101/2022.10.28.514214

**Authors:** EV Ho, A Welch, JA Knowles, SC Dulawa

## Abstract

Protein Tyrosine Phosphatase receptor type D (PTPRD) is a member of the protein tyrosine phosphatase family that mediates cell adhesion and synaptic specification. Genetic studies have linked *Ptprd* to several neuropsychiatric phenotypes, including Restless Leg Syndrome (RLS), opioid abuse disorder, and antipsychotic-induced weight gain. Genome-wide association studies (GWAS) of either pediatric obsessive-compulsive traits, or Obsessive-Compulsive Disorder (OCD), have identified loci near *Ptprd* as genome-wide significant, or strongly suggestive for this trait. We assessed *Ptprd* wild-type (WT), heterozygous (HT), and knockout (KO) mice for behavioral dimensions that are altered in OCD, including anxiety and exploration (open field test, dig test), perseverative behavior (splash-induced grooming, spatial *d*), sensorimotor gating (prepulse inhibition), and home cage goal-directed behavior (nest building). No effect of genotype was observed in any measure of the open field test, dig test, or splash test. However, *Ptprd* KO mice of both sexes showed impairments in nest building behavior. Finally, female, but not male, *Ptprd* KO mice showed deficits in prepulse inhibition, an operational measure of sensorimotor gating that is reduced in female, but not male, OCD patients. Our results indicate that constitutive lack of *Ptprd* may contribute to the development of certain domains that are altered OCD, including goal-directed behavior, and reduced sensorimotor gating specifically in females.

## Introduction

Obsessive-Compulsive Disorder (OCD) is a common and disabling neuropsychiatric disorder with a lifetime prevalence of approximately 2–3% (1). OCD is characterized by intrusive thoughts, images, or impulses, and/or repetitive compulsive behaviors (2). OCD is widely considered to be polygenic in origin (3-5), and family and twin studies have estimated heritability as high as 50% (6, 7). Two genome-wide association studies (GWAS) of OCD have been published to date, as well as a meta-analysis (8-10). Both the GWAS by Stewart et al. (2013) and the meta-analysis identified a variant located near the gene *protein tyrosine phosphatase, receptor type, D (Ptprd)* as one of the top signals (8), although neither study identified any single-nucleotide polymorphism (SNP) as genome-wide significant. Additionally, a GWAS of pediatric obsessive-compulsive traits identified a locus within in an intron of *Ptprd* that achieved genome-wide significance (11). Thus, although GWAS studies to date remain underpowered for identifying genome-wide significant single nucleotide polymorphisms (SNPs) for OCD or OC traits, *Ptprd* has emerged from these studies as a top candidate gene of interest.

Both common and rare variants have been identified for PTPRD, a cell-adhesion molecule and synaptic specifier which has been implicated in a number of brain phenotypes and disorders (12). For example, common variation in *Ptprd* has been associated with Restless Leg Syndrome (13, 14), which is characterized by irresistible urges to move the limbs and repetitive movements of the affected limb that serve to quench these urges (15, 16). Furthermore, SNPs in the *Ptprd* locus were associated with neurofibrillary tangle density in Alzheimer’s disease and accounted for 3% of individual differences in neurofibrillary pathology (12, 17). Other common SNPs in *Ptprd* have been implicated in atypical antipsychotic-induced weight gain (18, 19). A homozygous *Ptprd* micodeletion was identified in an individual with intellectual disability (20); however, more evidence will be required to establish a role for *Ptprd* deletion in cognitive disability. Thus, *Ptprd* plays a role in many brain-related phenotypes, in addition to OCD.

PTPRD undergoes homomeric binding and acts as a hemophilic, neurite-promoting cell adhesion molecule for neurons (21). In addition to homomeric binding, PTPRD also binds to additional ligands. For example, PTPRD has been reported to bind to interleukin 1 receptor accessory protein (22), netrin-G ligand-3 (NGL-3)(23), and synaptic adhesion-like molecules (SALMs)(24). Interestingly, PTPRD also directly binds with all SLIT and NTRK-like (SLITRK) family member proteins, including SLITRK1 and SLITRK5 (25, 26), which have been implicated in OCD by rodent preclinical work and human genetic studies evaluating rare variants (27-30).

To date, no animal studies to date have examined the role of *Ptprd* on behavioral phenotypes relevant to OCD. Here, we performed an initial behavioral characterization of *Ptprd* WT, HT, and KO mice of both sexes for behavioral constructs with relevance to OCD. We evaluated mice in the open field and the dig test to assess exploration and anxiety, which are reduced and increased in OCD, respectively (31-34). To assess perseverative behavior, we assessed route stereotypy in the open field using spatial *d* (35-37), and splash-induced grooming to assess perseverative grooming behavior (38, 39). To evaluate goal-directed behavior, which is reduced in OCD (40-42), we assessed nest building in the home cage (43-45). Finally, we measured prepulse inhibition (PPI) to assess sensorimotor gating (46-49), which is reduced preferentially in females, and not males, with OCD (50, 51).

## Methods

### Animals

*Ptprd* knockout (KO) mice on a mixed 129×1/SvJ x 129S1/Sy background were generously provided by Dr. Noriko Uetani at McGill University (52). Experimental cohorts were generated from heterozygous (HT) *Ptprd* crosses. Male and female wildtype (WT), heterozygous (HT), and knockout (KO) young adult mice 8-11 weeks of age were used for experiments (females: 16 WT, 16 HT, 9 KO; males: 16 WT, 16HT, 11 KO). Mice were housed under a 12hr light/dark cycle (lights on at 0700) with food and water available *ad libitum*. Behavioral testing was performed during the light-on portion of the light cycle. All methods and animal care procedures were prospectively approved by the University of California San Diego Animal Care and Use Committee (Protocol Number: S15266), and all aspects of the study were carried out in accordance with the recommendations of the National Institutes of Health’s Guide for the Care and Use of Laboratory Animals. At the end of experiments, mice were sacrificed using carbon dioxide inhalation using an Animal Care Panel approved apparatus followed by cervical dislocation.

### Open Field

Mice were placed in the corner of automated activity chambers, which were 41cm x 41cm with a 16 × 16 photobeam grid, and 2.54cm between each photobeam (Accusan, Columbus, OH). Activity was recorded for 60 minutes. Total distance traveled was quantified to assess locomotor activity. Vertical activity was used to assess rearing, an exploratory behavior. Percent center distance was calculated as [(center distance/total distance) x 100] and was used as a measure of anxiety, with increases in center distance interpreted as a decrease in anxiety. Spatial *d* was calculated using BMDP, Python and Night Owl to assess the degree to which consecutive movements were along a straight line (*d*∼1), meandering (*d*∼1.5), or contained many directional changes (*d*∼2).

### Dig Test

Immediately after open field, mice were placed in a standard cage with fresh bedding 1” deep and video recorded for 3 minutes. Videos were then scored by an experimenter blind to genotype and sex. Measures were latency to dig, total time digging, number of bouts digging, and average bout duration. A digging bout was defined as significant displacement of bedding due to limb or nose movement lasting at least one second, and not separated by more than one second of rest. Increases in digging behavior indicated increased exploration.

### Splash test

Following the dig test, mice were allowed to habituate to the test cage for 1 hour. Mice were then removed from the test cage, sprayed twice on their dorsal surface from approximately 5” away with a 10% sucrose solution, and returned to the test cage. Behavior was videotaped for 5 minutes, and then scored by a blind experimenter for latency to groom, total time grooming, number of grooming bouts, and average bout duration. A grooming bout was defined as any number of leg strokes along the body, a minimum of 2 arm strokes over the face/head, or any amount of time spent licking/biting the fur, with a break not separated by more than 2 seconds. Decreases in grooming in response to sucrose spray were interpreted as increased anhedonia.

### Nest Building

Nest building was used to assess general home cage behavior. Mice were placed into a novel cage with an unused nestlet (a 2” x 2” square of cotton fiber; Ancare). Nests were photographed at 0, 2, 4, and 6 hrs. Nests were scored by an experimenter blinded to genotype and sex, as described previously. Briefly, nests were given a score of 0-5 according to the following criteria: 0 = untouched, 1 = minimal pieces ripped off; 2 = major pieces of nestlet square left intact; 3 = all/most ripped up but not organized; 4 = all ripped up, organized into clear nest but not perfect, no tight walls; 5 = all ripped up, organized into clear nest, tight walls. Additionally, nests were weighed at time 0 and 6 hours, and the amount of nestlet used was calculated by subtracting the intact portion of the nestlet remaining at 6 hours from the original nestlet weight at time 0. Reductions in nest building are thought to reflect reduced well-being (53, 54).

### Prepulse Inhibition

Prepulse inhibition was used as an index of sensorimotor gating, which is reduced in female patients with OCD (50). Mice were placed into startle chambers that consisted of a Plexiglas cylinder resting on a Plexiglas platform in a sound-attenuating, ventilated chamber (San Diego Instruments, CA). Sessions were 20 minutes long. The first 5 minutes consisted of acclimation to the background (65 dB) noise, followed by four consecutive blocks of test trials. Testing consisted of a pseudo random representation of five different types of trials: a 40-millisecond broadband 120 dB burst (pulse alone trial); three different prepulse pulse trials in which 20-millisecond long 3 dB, 6 dB, or 12 dB above background stimuli preceded the 120 dB pulse by 100 milliseconds (onset to onset); and a no stimulus trial, in which only background noise (65 dB) was presented. Blocks one and four consisted of six consecutive Pulse alone trials, while blocks two and three contained six Pulse alone trials, five pp3p120 trials, five pp6p120 trials, five p12p120 trials and four No stimulus trials. Thus, the entire session consisted of 62 test trials. PPI was calculated as [(startled amplitude pulse – startle amplitude prepulse + pulse)/startle amplitude pulse) x 100]. Reduced PPI is thought to reflect impaired sensorimotor gating. Startle magnitude was calculated as the average response to 120 dB pulse trials from blocks two and three. Startle habituation was assessed as the average of 120 dB pulse trials from all 4 blocks.

### Statistical Analysis

Multi-way ANOVAs were performed for all measures with sex and genotype as between-subjects factors. Main effects of genotype were resolved using Student Newman-Keuls post hoc tests. For the analysis of the open field, nest building test, and startle habituation, time was an additional within subjects factor. For the analysis of PPI, both prepulse intensity and block were additional within-subjects factors. Interactions were resolved using Student Newman-Keuls post hoc tests or post hoc ANOVAs with the Bonferonni correction applied as appropriate. For all measures, outliers were removed if their value was greater than two standard deviations above or below the mean. Alpha was set at 0.05.

## Results

### Open Field

The measures of total distance traveled (Figure 1a), center time (Figure 1b), center distance traveled (Figure 1c), percent of total traveled time spent in center (Figure 1d), rearing (Figure 1e), and spatial *d* (Figure 1f) were not altered by *Ptprd* genotype or sex, and no interactions were observed.

**Figure 1.**
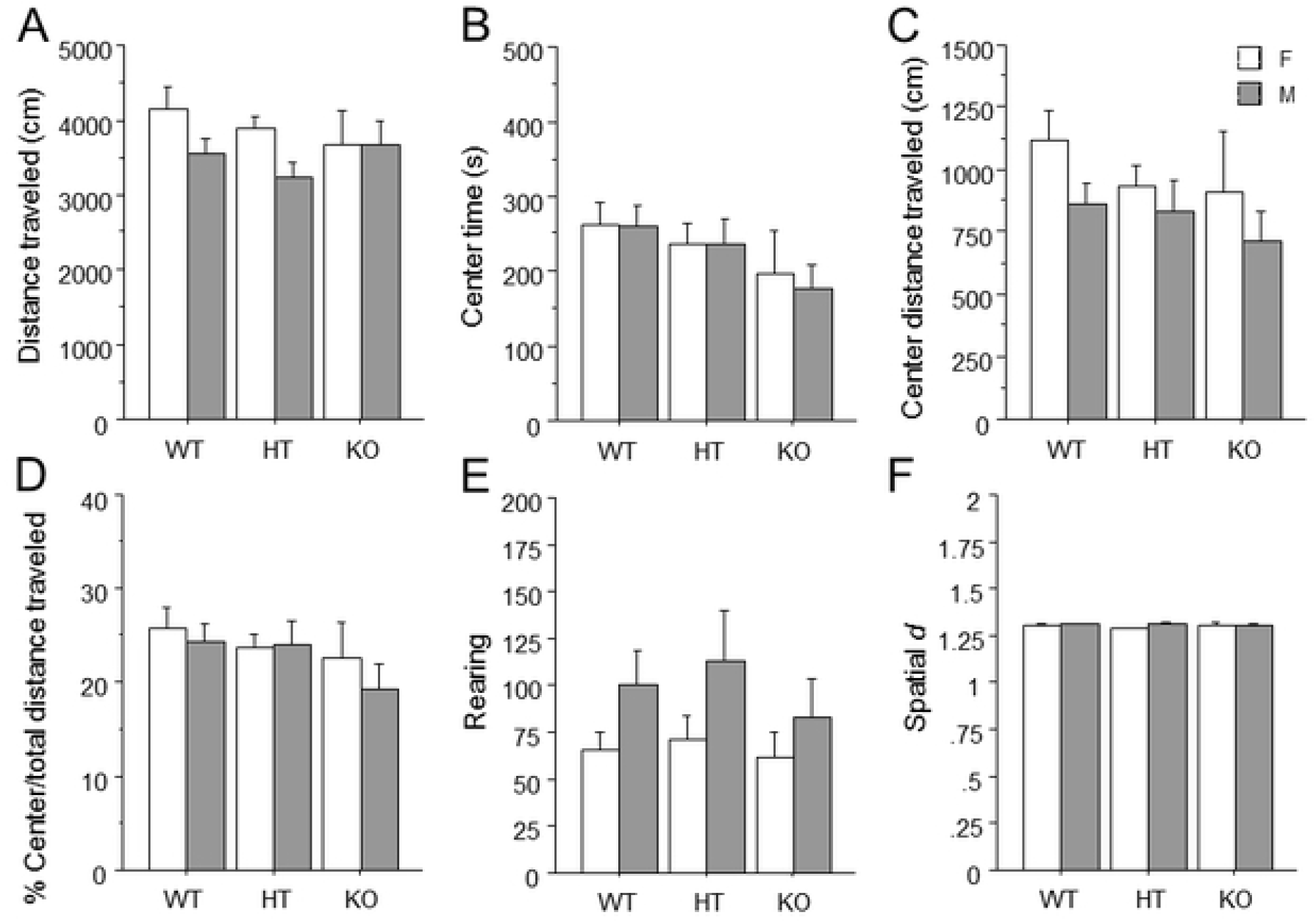
Unaltered open field behavior in mice lacking PTPRD. (A) Total distance traveled. (B) Total time spent in the center. (C) Total distance traveled in the center. (D) Percent distance traveled in the center divided by total distance traveled. (E) Total number of rearings. (F) The average spatial *d* value over the entire session. Mean ± s.e.m, n = females: 16 WT, 16 HT, 9 KO; males: 16 WT, 16HT, 11 KO.

### Dig Test

Time spent digging (Figure 2a) and the measures of latency to dig (Figure 2b), number of dig bouts (Figure 2c), and average duration of dig bout (Figure 2d) were not altered by genotype or sex. No interaction was observed for time spent digging [F(1,2) = 2.00, p = .14].

**Figure 2.**
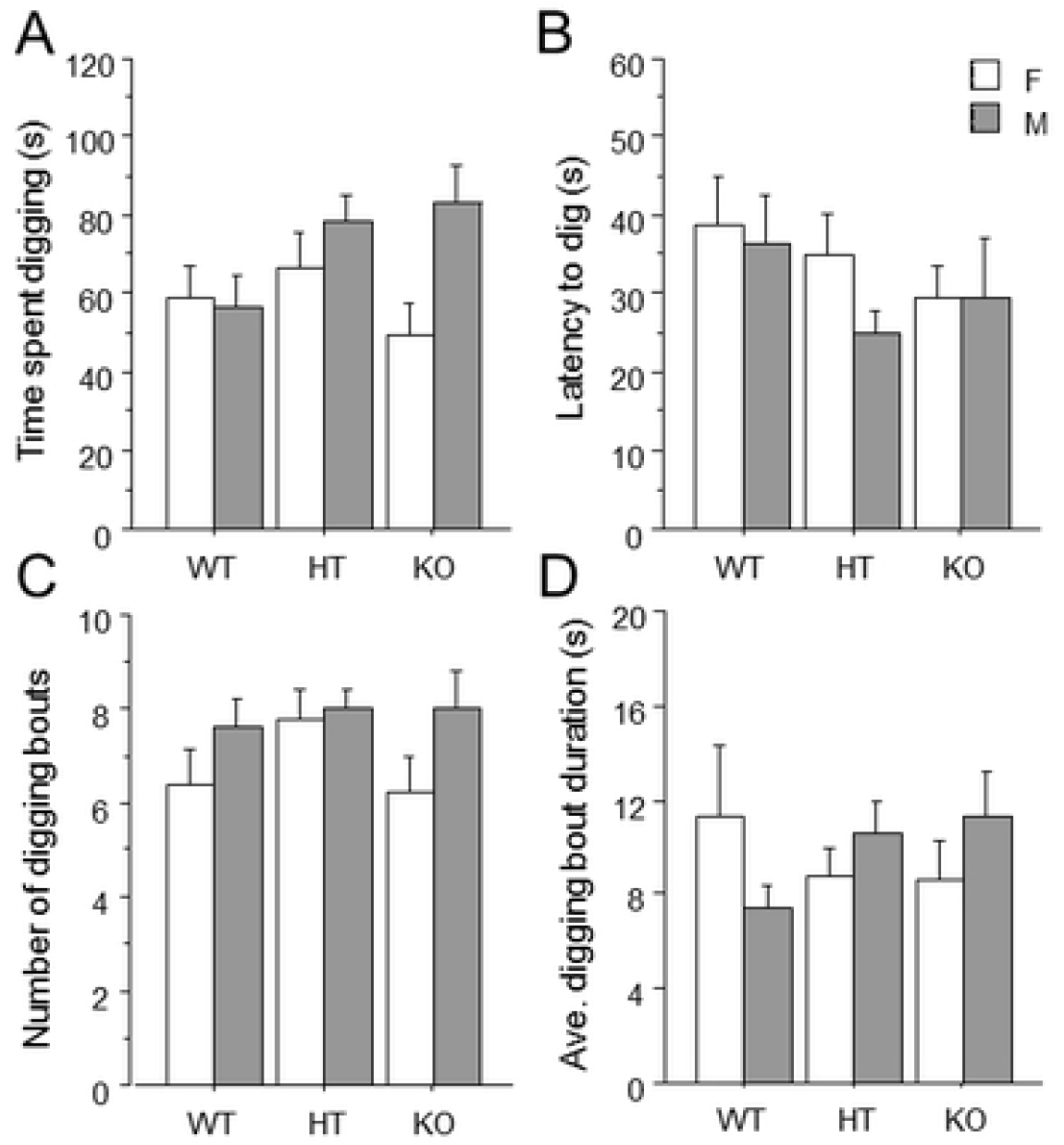
Unaltered digging behavior in mice lacking PTPRD. (A) Total time spent digging. (B) Latency to dig. (C) Total number of digging bouts. (D) The average duration of all digging bouts. Mean ± s.e.m.

### Splash Test

The measures of time spent grooming (Figure 3a), latency to groom (Figure 3b), number of grooming bouts (Figure 3c), and average groom bout duration (Figure 3d) were not altered by *Ptprd* genotype or sex.

**Figure 3.**
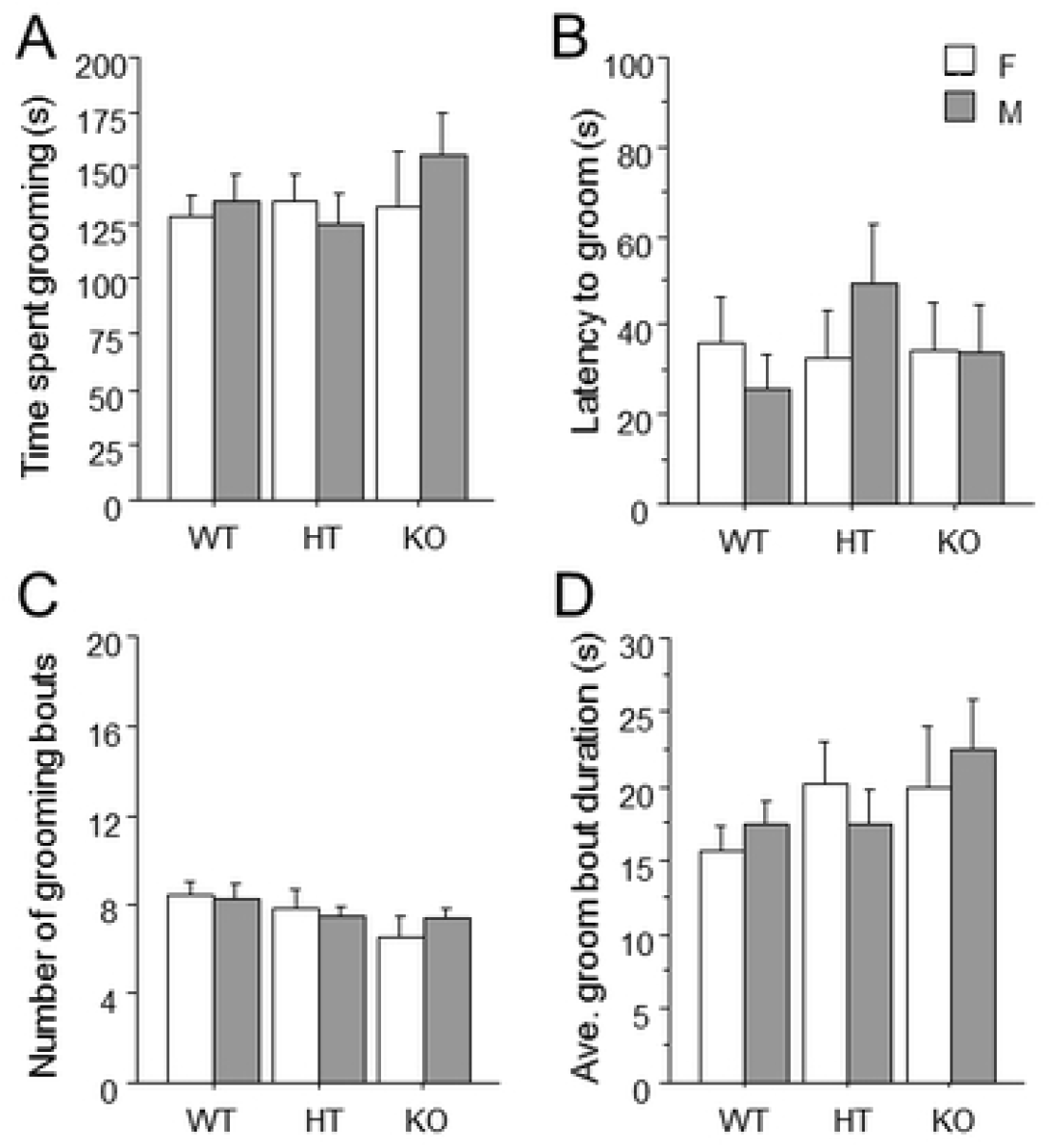
Unaltered plash-induced grooming behavior in mice lacking PTPRD. (A) Total time spent grooming. (B) Latency to groom. (C) Total number of grooming bouts. (D) The average duration of all grooming bouts. Mean ± s.e.m.

### Nest Building

ANOVA revealed that genotype [F(2, 78) = 8.34] (p<0.05) influenced Nestlet score (Figure 4a), and an interaction of genotype and block [F(8, 312) = 2.25] (p<0.05) was observed. Post-hoc analysis revealed that *Ptprd* KO mice made worse nests at the 2, 4, and 6 hour timepoints compared to *Ptprd* WT and HT mice. Additionally, genotype [F(2, 78) = 3.618] (p<0.05) influenced the percent of nestlet used (Figure 4b), with *Ptprd* KO mice using less percent nestlet than *Ptprd* HT or WT mice.

**Figure 4.**
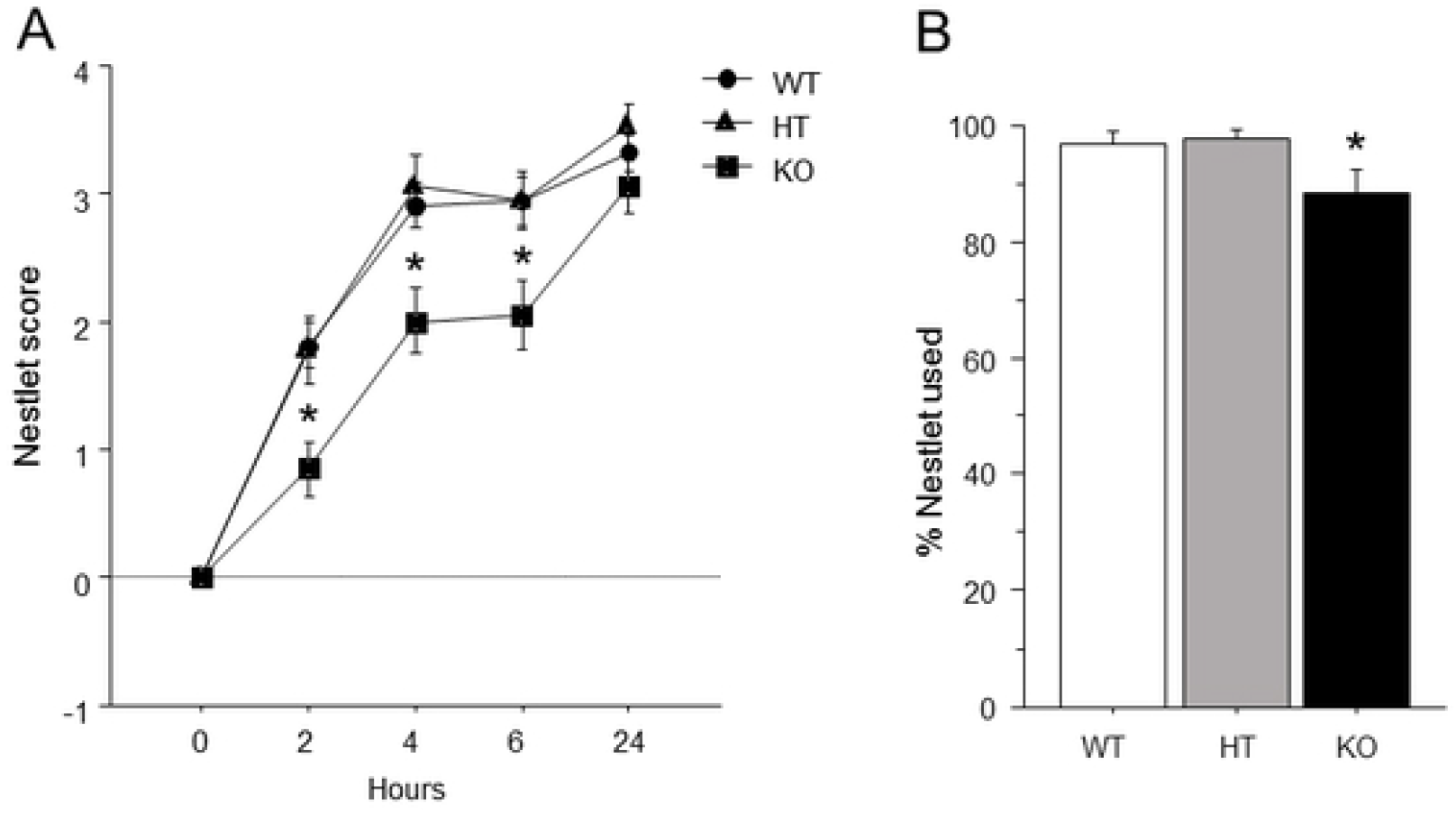
Impaired nest-building behavior in mice lacking PTPRD. (A) Nestlet score at 0, 2, 4, 6, and 24 hours after the beginning of the test. (B) Percent of the nestlet used by the end of the test. Mean ± s.e.m. An asterisk indicates a significant difference from both HT and WT mice.

### PPI

ANOVA revealed a genotype by sex interaction [F(2, 78) = 3.31] (p<0.05) for percent prepulse inhibition; no other interactions were found. Post-hoc ANOVAs examining effects of *Ptprd* genotype within each sex identified an effect of *Ptprd* genotype within females [F(2, 38) = 3.58] (p = .038)(Figure 5a), but not males. However, this p value fell just short of the p = 0.025 cut-off following Bonferroni correction. We performed Newman-Keuls planned comparisons, which revealed that within females, *Ptprd* KO mice showed reduced percent prepulse inhibition relative to both *Ptprd* WT and HT mice. Additionally, post-hoc Newman-Keuls revealed that within *Ptprd* knockout mice, females had lower percent prepulse inhibition than males.

**Figure 5.**
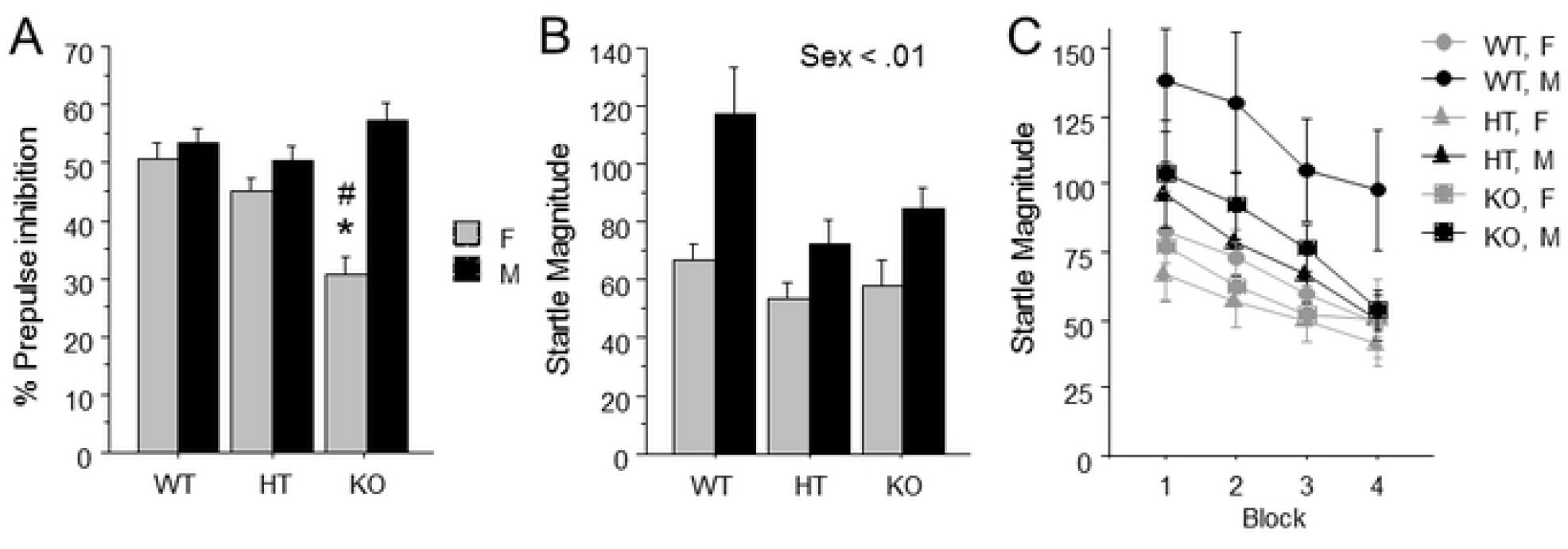
Reduced percent prepulse inhibition only in female mice lacking PTPRD. (A) Percent prepulse inhibition averaged across prepulse intensity. (B) Average startle magnitude during blocks two and three. (C) Average startle magnitude is shown for each of four blocks. Mean ± s.e.m. An asterisk indicates a significant difference from both HT and WT mice of the same sex. A pound sign indicates a significant difference from males within the same genotype.

ANOVA revealed a main effect of sex [F(1, 78) = 8.51] (p<0.01) on startle magnitude, with males showing a larger average startle magnitude than females (Figure 5b). Additionally, an ANOVA assessing the effects of sex and genotype across all four blocks also found a main effect of sex [F(1, 78) = 8.78] (p < 0.01), with males showing larger startle magnitudes than females. However, no interactions including block were observed, indicating there were no differences in startle habituation between any of the groups (Figure 5c).

## Discussion

Here we show that *Ptprd* KO mice show impairments in nest building behavior, a goal directed behavior performed by mice in the home cage, and deficits in PPI only in females. The other behavioral constructs that were assessed including anxiety/exploration, perseverative behaviors, and locomotion, were unaltered by *Ptprd* genotype. Specifically, we did not observe any anxiety-related changes in center measures of the open field, or changes in overall locomotion between genotypes. We also did not find any effect of *Ptprd* genotype on exploratory measures in the open field including rearing behavior, or changes in the dig test. Finally, *Ptprd* genotype did not affect the expression of perseverative behaviors including splash-induced grooming or route stereotypy. Thus, *Ptprd* KO mice showed deficits in two constructs that are impaired in OCD patients, which are goal-directed behavior and sensorimotor gating. However, showing deficits in these two behavioral domains is not specific to OCD, but is also characteristic of other neuropsychiatric disorders including schizophrenia (55-59). Although variation in *Ptprd* has been linked to OCD, no significant association has yet been made to schizophrenia by human genetic studies.

We evaluated mice in the open field and the dig test to assess exploration and anxiety, since patients with OCD show increases in anxiety and reductions in exploration or novelty-seeking behavior (31-34, 60, 61). However, we observed no effect of *Ptprd* genotype on measures of exploration in the open field including rearing (Figure 1e), or on digging measures of the dig test (Figure 2). Furthermore, we found no effect of *Ptprd* genotype on anxiety-related measures of center time, center distance, or center/total distance traveled in the open field (Figure 1b-d). It should be noted that OCD was removed from the category of anxiety disorders in DSM V, and was placed under the heading of ‘Obsessive-Compulsive and Related Disorders’. The rationale for this change was that obsessions and compulsions, and not anxiety, are the fundamental features of the disorder. For example, OCD patients often experience negative affect including disgust, discomfort, and unease in relation to particular stimuli, but not all patients experience anxiety (62, 63). Therefore, while the open field test provides a classical measure of anxiety, this measure might not capture other forms of negative emotions. For example, center measures of the open field are highly responsive to classical anxiolytic drugs (64), while the obsessions and compulsions of OCD are not (65, 66). We did not find any effect of *Ptprd* genotype on overall locomotor activity in the open field (Figure 1a). These findings are in contrast to a previous report suggesting that *Ptprd* HT mice show small increases in locomotion in a conditioned place preference apparatus compared to *Ptprd* WT and KO mice (67). In addition to finding comparable locomotion in all genotypes, we did not notice any hind limb weakness in *Ptprd* KO mice, which was previously reported (52).

Grooming behavior has been frequently assessed in rodents in an attempt to study the repetitive behaviors observed in OCD patients; however, this approach relies largely on face validity. Indeed, several mouse genetic models that show robust overgrooming behavior, including SAPAP3 KO and SLTRK5 KO mice (29, 68-70) are constitutive knockout mouse models for genes that have been tentatively linked with OCD (28, 71). However, a definitive link for these genes with OCD remains to be established by large scale human genetic studies. We did not observe any effect of *Ptprd* genotype on grooming behavior in the splash test (Figure 3), which involves spraying the dorsal surface of the mouse to elicit grooming behavior. We also investigated the role of *Ptprd* in another form of perseverative behavior, which is route stereotypy in the open field. Route stereotypy refers to repetitive circling of the edges of the open field along with increased smoothness of the locomotor path, and has been observed other mouse studies of OCD-related behavior (35, 36, 38, 72). Since we did not find any changes in spatial *d* (Figure 1f), which quantifies the smoothness of the path of the animal, or reductions in center measures in the open field, *Ptprd* genotype did not alter route stereotypy. In summary, we did not find any effect *Ptprd* genotype on perseverative behaviors in the tests employed.

OCD patients exhibit reductions in goal-directed behavior and increases in habitual behavior (40-42). Indeed, compulsive behaviors are defined as maladaptive repetitive behaviors that are performed despite no longer leading to a goal, and are typically inflexible in nature (73-75). Rodent nest building is considered a goal-directed behavior (43-45) that can be readily assessed in the home cage. We previously reported that mice lacking *Btbd3*, another top OCD candidate gene identified in the GWAS by Stewart et al. (2013), show reductions in nestlet behavior (36). Here, we found that *Ptprd* KO mice had lower nest building scores than *Ptprd* WT and HT mice at the 2, 4, and 6 hour timepoints (Figure 4a). Although they achieved the same nestlet score as the other genotypes by 24 hours, they also used less of the nestlet overall compared to *Ptprd* WT and HT mice (Figure 4b). Future studies will evaluate *Ptprd* KO mice in a probabilistic learning task (PLT) which can be used to measure goal-directed versus habitual decision-making strategies in both rodent and humans (76, 77), with OCD patients showing a shift towards habitual choices in this paradigm (78, 79).

PPI is a form of startle plasticity that can be measured similarly in both humans and animals, and provides an operational measure of sensorimotor gating. PPI is reduced in OCD patients (51, 80, 81), and intrusive thoughts, images and impulses and repetitive actions are thought to reflect failure to gate incoming stimuli and outgoing motor routines, respectively (81). A more recent study assessing PPI in OCD patients using more subjects suggested that among OCD patients, the deficits in PPI are more pronounced in females than males (50). Similarly, we found that female, but not male, *Ptprd* KO mice showed reductions in PPI compared to female *Ptprd* WT and HT mice, as well as male *Ptprd* KO mice (Figure 5a). The reduction in PPI observed in female *Ptprd* KO mice was found across all prepulse intensities, as with female OCD patients reported in Ahmari et al. (2016). The reduction in PPI observed in female *Ptprd* KO mice was not an artifact of any effect of *Ptprd* genotype on block 2 and 3 startle magnitude, which is used in the calculation of PPI values. Specifically, no effect of *Ptprd* genotype was found on block 2 and 3 startle values (Figure 5b); only a main effect for males to show higher startle overall was observed. There was also no effect of *Ptprd* genotype on startle habituation across blocks 1-4 (Figure 5c). Consistent with this finding, patients with OCD exhibit startle habituation that is comparable to control subjects (51, 80). However, patients with other neuropsychiatric disorders, such as schizophrenia, exhibit deficits in both PPI and startle habituation (59, 82). In summary, *Ptprd* KO mice show a sexually dimorphic reduction in PPI without any change in startle habituation, similar to OCD patients.

Whether PTPRD function is increased or decreased in OCD patients or individuals with OC-traits remains unresolved. Although another study reported one case of a copy number variant (CNV) duplication including *Ptprd* (83) in pediatric OCD, more work will be required to determine whether increase or decrease of *Ptprd* function is linked with OCD. To our knowledge, only *Ptprd* KO mice, and not *Ptprd* overexpressing mice, are available for study. Here, we found that *Ptprd* KO, but not *Ptprd* HT, mice showed two phenotypes that are exhibited by OCD patients: reduced goal-directed behavior, and reduced PPI in females only. However, our findings do not rule out the possibility that *Ptprd* KO mice might show increases in perseverative behavior or anxiety in other behavioral tests. Future studies should investigate the behavioral consequences of overexpressing *Ptprd* in mice to further clarify the role of *Ptprd* function in OCD-related behavioral domains. Furthermore, additional information regarding the status of *Ptprd* function in OCD patients would permit the generation of more useful animal genetic models.

Together, the results presented here show that loss of *Ptprd* function reduces goal-directed behavior in the home cage, and disrupts PPI specifically in females, which are two phenotypes that are directly relevant to OCD. Future studies should further confirm the role of *Ptprd* in goal-directed behavior using a translational behavioral paradigm that has yielded differences between OCD patients and controls. Furthermore, future studies should investigate the role of gain-of-function of *Ptprd* to further clarify the role of this gene in regulating OCD-relevant domains. Such animal studies and more human genetic studies will be critical to determine the role of *Ptprd* variation in risk for OCD and OC-traits.

## Notes

Funding: This work was supported by the National Institute of Mental Health (NIMH) (https://www.nimh.nih.gov/) grant R21-MH128574 (S.C.D.), and the Della Martin Foundation (J.A.K.).

### Competing Interest Statement

The authors have declared no competing interest.

